# Markerless mouse tracking for social experiments

**DOI:** 10.1101/2021.10.20.464614

**Authors:** Van Anh Le, Toni-Lee Sterley, Ning Cheng, Jaideep S. Bains, Kartikeya Murari

**Affiliations:** Electrical and Software Engineering, University of Calgary; Hotchkiss Brain Institute, University of Calgary; Department of Comparative Biology and Experimental Medicine, Faculty of Veterinary Medicine, University of Calgary; Alberta Children’s Hospital Research Institute, University of Calgary

## Abstract

Automated behavior quantification requires accurate tracking of animals. Simultaneous tracking of multiple animals, particularly those lacking visual identifiers, is particularly challenging. Problems of mistaken identities and lost information on key anatomical features are common in existing methods. Here we propose a markerless video-based tool to simultaneously track two socially interacting mice of the same appearance. It incorporates conventional handcrafted tracking and deep learning based techniques, which are trained on a small number of labeled images from a very basic, uncluttered experimental setup. The output consists of body masks and coordinates of the snout and tail-base for each mouse. The method was tested on a series of cross-setup videos recorded under commonly used experimental conditions including bedding in the cage and fiberoptic or headstage implants on the mice. Results obtained without any human intervention showed the effectiveness of the proposed approach, evidenced by a near elimination of identities switches and a 10% improvement in tracking accuracy over a pure deep-learning-based keypoint tracking approach trained on the same data. Finally, we demonstrated an application of this approach in studies of social behaviour of mice, by using it to quantify and compare interactions between pairs of mice in which some are anosmic, i.e. unable to smell. Our results indicated loss of olfaction impaired typical snout-directed social recognition behaviors of mice, while non-snout-directed social behaviours were enhanced. Together, these results suggest that the hybrid approach could be valuable for studying group behaviors in rodents, such as social interactions.

## 1 Background

Social interactions are a cornerstone of human society. Social deficits are a hallmark of several neurodevelopmental disorders such as autism spectrum disorders (DiCicco-Bloom et al., 2006) and attention deficit hyperactivity disorder (Harpin et al., 2016). Animal models, particularly mouse models, are widely used to investigate social behavior and the mechanisms of such diseases. Measurements of social interactions in mice under normal, diseased and treatment conditions are important components of such studies. The assessment of social behavior, commonly conducted by human annotators, can be subjective, time-consuming, and labor-intensive. There is increasing interest in using automated tools which can overcome these limitations and allow accurate and high throughput analysis of animal social behavior.

Several automated analysis tools are based on an estimation of the posture and position of animals. Therefore, the quality of tracking algorithms has a large impact on assessment accuracy (Lorbach et al., 2015). Visual object tracking is a fundamental and crucial problem in machine vision with several real-world applications (Smeulders et al., 2013; Yilmaz et al., 2006). Despite rapid progress in computer vision, tracking animals in their natural environment is still a daunting task due to factors like illumination changes, background clutter, occlusion from vegetation and other natural features, shape deformation and abrupt motion (Dell et al., 2014). Within controlled laboratory settings, some of those challenges can be alleviated.

Currently, several methods exist for tracking animal locations in laboratory settings, and some have the added benefit of tracking key anatomical features. Approaches that use radio-frequency identification (RFID) (Galsworthy et al., 2005; Lewejohann et al., 2009; Schaefer and Claridge-Chang, 2012), can maintain individual identity well, and thus provide accurate tracking of positions of multiple animals. However, they do not provide information on the positions of important body regions. Techniques combining RFID and video tracking (Weissbrod et al., 2013; de Chaumont et al., 2019) or video tracking visually marked animals (Ohayon et al., 2013; Lorbach et al., 2017) offer both tracking and keypoint locations. However, animals need to go through additional preparations for these methods, adding cost of time and potential confound factors such as discomfort or inflammation. There are also concerns that these methods could alter animal behaviour (Walker et al., 2013; Burn et al., 2008; Hurst and West, 2010; Hershey et al., 2018).

Deep learning based methods can track animal positions, as well as extract key anatomical points such as the nose and tail-base, which are useful for further behavioural analysis. Inspired by the success of Convolutional Neural Networks (CNNs) in many computer vision tasks (Simonyan and Zisserman, 2014; He et al., 2016; Krizhevsky et al., 2012), several approaches have been proposed to utilize hierarchical features learned from CNNs for visual tracking (Qi et al., 2016; Danelljan et al., 2015; Yang and Chan, 2017; Fan et al., 2010; Wang and Yeung, 2013; Ma et al., 2015). Recently, such techniques have been applied to animal pose estimation (Mathis et al., 2018; Pereira et al., 2019; Graving et al., 2019). Furthermore, the latest version of DeepLabCut (Nath et al., 2019) allows tracking features of multiple identical-looking animals, and Social LEAP Estimates Animal Pose (SLEAP) (https://sleap.ai/, accessed November 17, 2021) provides a framework for multi-animal body part position estimation. Applying these frameworks on large datasets often requires iterative refinement using a large training set and human correction in post-processing. This is due to swapped or missed features and identities caused by visual occlusion during social interactions. In addition, there is a dearth of publicly available datasets for developing new techniques and comparing different approaches.

In this paper, we propose a markerless approach which combines a conventional tracking method and segmentation based on deep learning for tracking the identities of two interacting mice of the same appearance. Specifically, we use Mask R-CNN (He et al., 2017), a flexible and general framework for object instance segmentation, to segment the mice when they interact with each other. To train the model effectively, we propose a technique to capture a wide variety of mouse postures, while minimizing the cost of preparing training data. We also track key points including snout and tail-base, which are important for quantifying distinct social behaviors. Furthermore, we incorporate a hand-designed module to assist deep learning based pose estimation approaches, such as DeepLabCut or SLEAP, to address issues such as lost key points due to occlusion, which are common when the animals are in close contact.

To validate the proposed system and test the extent of generalization, we trained the algorithm using images from a simple, clutter-free setup. We then applied it to videos collected in different relevant setups with a wide range of confounding variables such as bedding and fiberoptic or electrophysiology implants. The results were then evaluated using previously established metrics and compared with the state-of-the-art automated methods and human annotation. Results indicated that our proposed approach is effective; specifically, we reduced snout and tail-base location errors and the number of identity switches by 1-2 orders of magnitude, and improved multi object tracking accuracy by over 10% compared to using DeepLabCut alone.

To demonstrate the utility of this approach, we used our system to address a biological question. It is known that mice and rats typically show a preference for social novelty, spending more time investigating an unfamiliar versus a familiar conspecific (Moy et al., 2004; Oettl et al., 2016). To discriminate between familiar and unfamiliar conspecifics, rodents rely on olfactory cues (Sanchez-Andrade and Kendrick, 2009; Oettl et al., 2016). Here we asked how social discrimination of a familiar versus an unfamiliar conspecific will be affected by loss of olfaction, i.e. anosmia. We used intranasal ZnSO_4_ to induce anosmia in some animals (Ducray et al., 2002), while control animals received intranasal saline. We then assessed how anosmic and control mice behaved towards familiar or unfamiliar conspecifics. From the automated analyses we were able to make several conclusions regarding how anosmia impacts mouse social behaviors towards familiar and unfamiliar conspecifics.

## 2 Experimental setups and Mouse Tracking dataset

Experiments were conducted in 12 different settings as shown in Table 1. Figure 1 shows sample images from the videos highlighting the different settings. In all cases, the animals were placed in a cage measuring 30 × 30 × 30 cm^3^ with the walls lined with a diffuse material. A camera was installed on the lid of the cage and pointed towards the bottom of the cage. Infrared lights on the cage lid and an infrared bandpass filter on the camera were used to ensure constant image brightness regardless of ambient lighting and maintain good contrast. All the videos were recorded at 30 frames per second with a resolution of 540 × 540 pixels. A total of 14 C57BL/6J, BTBR or Crh-IRES-Cre::Ai14 mice aged 7 to 20 weeks were used for the experiments. Mice were housed in standard mouse cages with access to food and water ad libitum, in a humidity- and temperature-controlled room with a 12-h light/dark cycle. All procedures for this study were performed according to the recommendations by the Canadian Council for Animal Care. The protocol of this study was approved by the Health Sciences Animal Care Committee of the University of Calgary.

**Figure 1:**
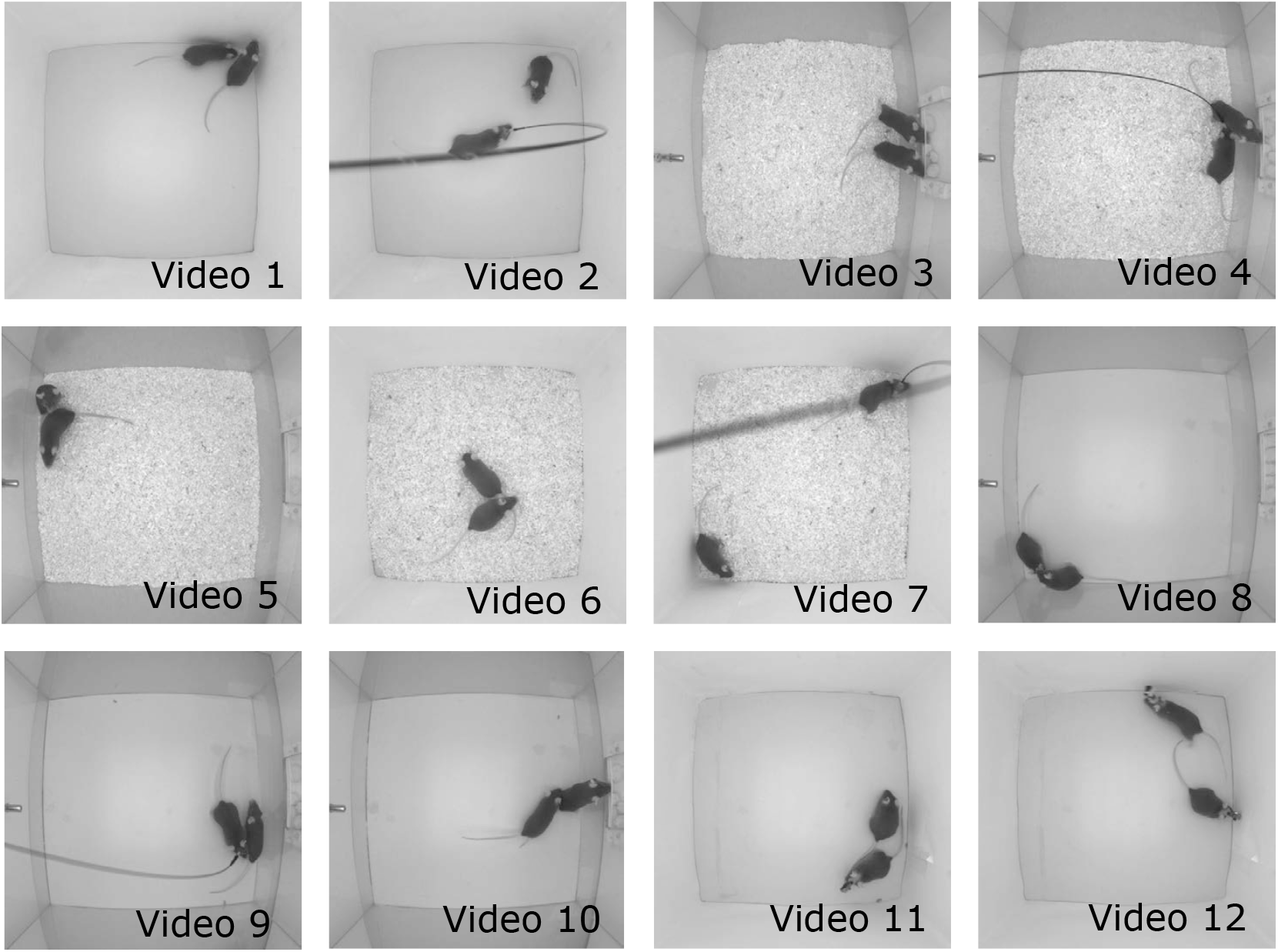
Snapshots taken from the videos illustrating twelve experimental setups used in the Mouse Tracking Dataset. Details for each of the settings are in Table 1.

**Table 1:** Details of the experimental settings used in the Mouse Tracking Dataset

| Video | Animal genotype | Sex | Bedding | Food and water | Implant |
| --- | --- | --- | --- | --- | --- |
| 1 | C57BL/6, C57BL/6 | f/f | - | - | -/- |
| 2 | Crh-IRES-Cre::Ai14, C57BL/6 | f/f | - | - | fiberoptic/- |
| 3 | C57BL/6, C57BL/6 | f/f | yes | - | -/- |
| 4 | Crh-IRES-Cre::Ai14, C57BL/6 | f/f | yes | yes | fiberoptic/- |
| 5 | C57BL/6, C57BL/6 | m/f | yes | yes | -/- |
| 6 | BTBR, BTBR | m/f | yes | - | -/- |
| 7 | Crh-IRES-Cre::Ai14, C57BL/6 | f/f | yes | - | fiberoptic/- |
| 8 | C57BL/6, C57BL/6 | f/f | - | yes | -/- |
| 9 | Crh-IRES-Cre::Ai14, C57BL/6 | f/f | - | yes | fiberoptic/- |
| 10 | C57BL/6, C57BL/6 | m/f | - | yes | -/- |
| 11 | C57BL/6, C57BL/6 | m/f | - | - | EEG/- |
| 12 | C57BL/6, C57BL/6 | m/m | - | - | EEG/EEG |

Four animals were implanted with fiberoptic or electrophysiology headstages. Twelve videos, each 10 minutes long, were obtained. We term this group of videos as the Mouse Tracking dataset (MT). For analysis, videos were grouped into four categories based on commonly encountered experimental settings - simple (videos 1, 8, 10), with bedding (videos 3-7), with tether (videos 2, 4, 7, 9) and with a headstage (videos 11, 12).

## 3 Methodology

### 3.1 Overview of the proposed approach

Figure 2 shows the top-down approach of our proposed algorithm. The input to the algorithm is a sequence of video frames which first go through a stage of conventional foreground detection. In this stage, a background model is created, either using images from before the experiment started or using a temporal median filter on the very first frames after the experiment starts. The foreground is obtained by thresholding with a threshold model which is chosen to be half of background intensity, followed by morphological operations (Supplementary Figure S1). Subsequently, the frames where the number of foregrounds is equal to the number of animals bypass the next step. In frames where this criterion is not satisfied, the detection is considered to have failed. This often occurs when the animals are in close physical proximity. These frames are sent to the Mask R-CNN model to segment the mice. The areas of output blobs generated by the Mask R-CNN are then verified to filter out segmentation errors in the post processing stage. In the next stage, tracking inference, the foreground masks of all frames are matched to subsequent frames to obtain the trajectories of each animal.

**Figure 2:**
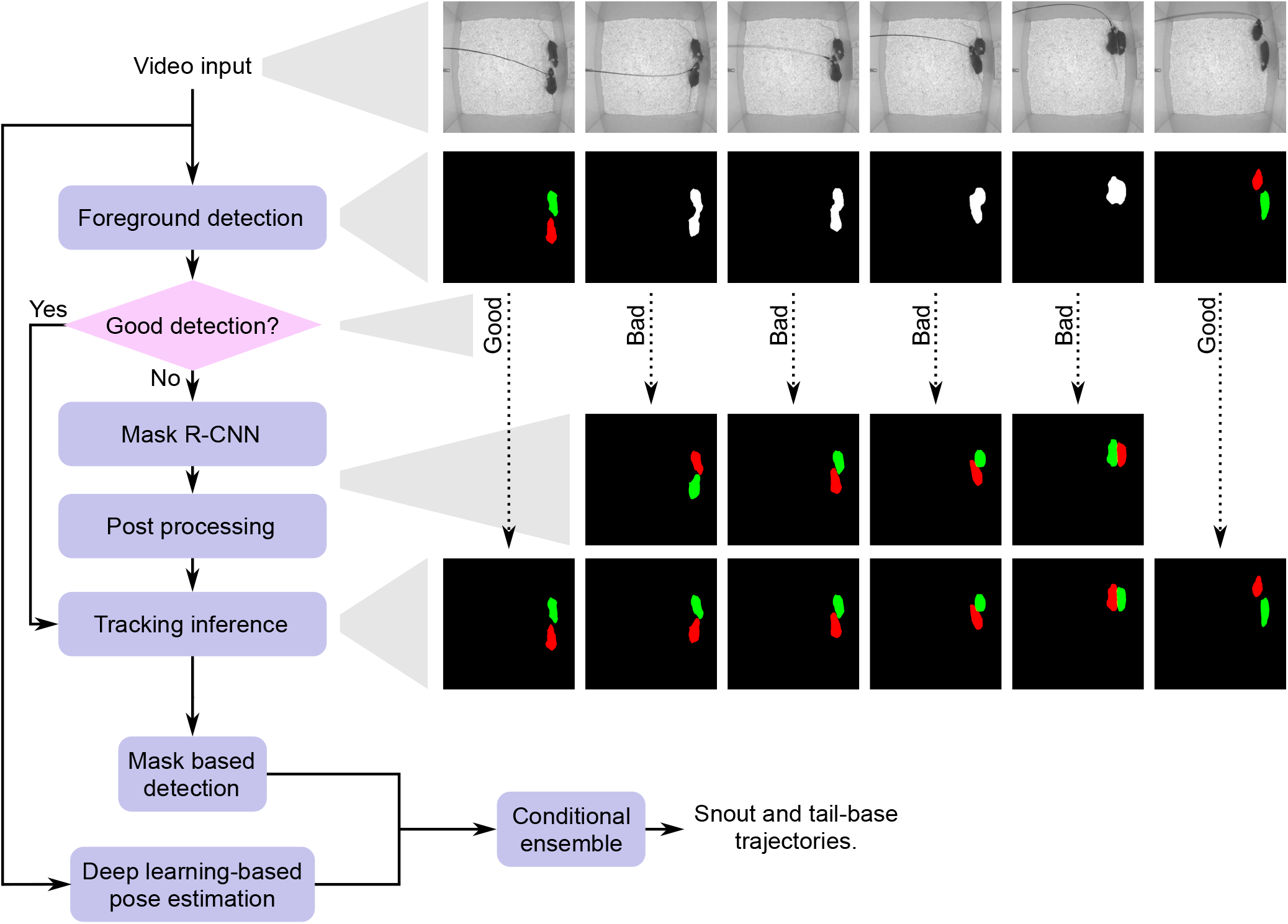
The pipeline of the proposed algorithm for mouse tracking and feature detection

Finally, key points consisting of snout and tail-base are detected via an ensemble stage combining detection results obtained by a hand-crafted approach termed mask-based detection (MD) and DeepLab-Cut. While any deep-learning-based pose estimation method can be used, we chose DeepLabCut based on its better performance over other methods Graving et al. (2019). In the following sections, we elaborate on the details of key modules and evaluation.

### 3.2 Data preparation

One drawback of deep-learning-based approaches (Nath et al., 2019; https://sleap.ai/, accessed November 17, 2021), is that extensive iterative training is required to reach a desired level of accuracy. In contrast, we aimed to obtain results with only one training session for each model and to investigate the limit of generalization of the approach when dealing with experimental variations. We generated training sets from a separate 7 minute video recorded in the most minimalistic setting *i.e.* with two C57BL/6 mice with no implants without bedding or any other enrichments. Since there are no confounders in this experimental setting, it achieves the best foreground separation through conventional background subtraction. Two foregrounds were obtained, i.e. both mice were successfully segmented, in about 84% of the frames. We randomly selected 10% of these to create Dataset 1 (1060 frames). It should be noted that Dataset 1 is obtained with no human intervention. In addition, we randomly chose ~10% of the video frames (212 frames) where the conventional foreground detection algorithm failed to separate mice due to physical proximity. We then manually segmented those frames using the Labelme annotation tool (Wada, accessed November 17, 2021) to build Dataset 2. Several images taken from Dataset 1 and 2 are shown in Fig. 3a and b. Both Dataset 1 and Dataset 2 were used to train the Mask R-CNN. In addition, Dataset 2 was used to create training data for DLC, as instructed in the documentation. Although our approach outputs the coordinates of only two key points (snout and tail-base), we over-labeled the training sets to obtain the best results. Particularly, each image was annotated with 12 points: snout, left ear, right ear, shoulder, spine 1, spine 2, spine 3, spine 4, tail-base, tail 1, tail 2, and tail end.

**Figure 3:**
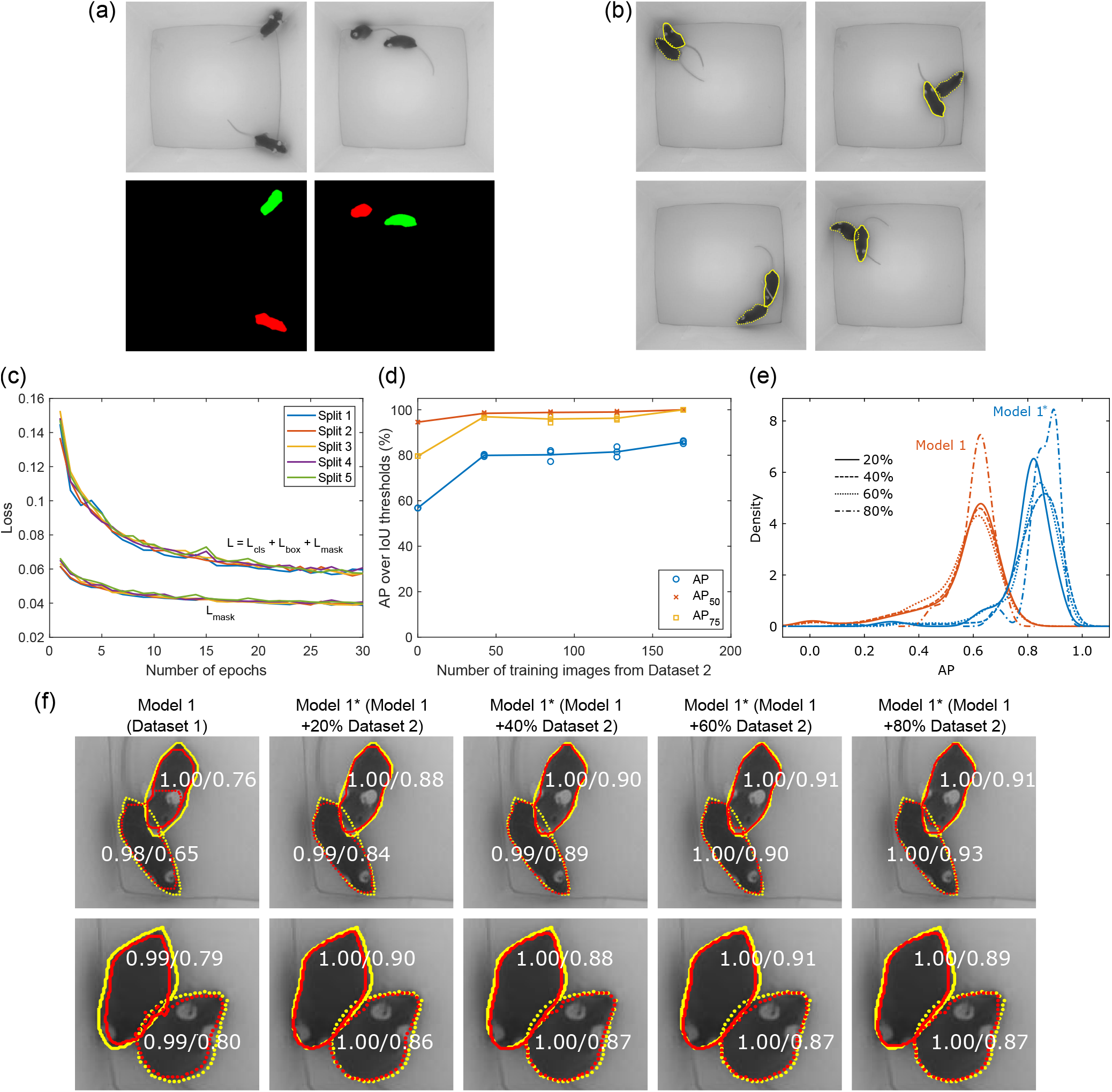
Training and evaluation of the Mask R-CNN. (a) Example images with labels taken from Dataset 1. (b) Example images with human annotations taken from Dataset 2. (c) Loss for 5-fold crossvalidation on Dataset 1. (d) *AP*, *AP*_75_ and *AP*_50_ of test data for 3 splits of training data vs. number of training images corresponding to 0, 20, 40, 60, and 80% of Dataset 2. (e) Kernel density estimate of *AP* of Model 1 and Model 1*s on test data for the first split of training set which account for 20-80% of Dataset 2. (f) Visualization of the outputs of Model 1 and Model 1*s with 20, 40, 60, and 80% training fraction in Dataset 2 (red) and human annotation (yellow). Each row shows performance on a different frame. Number pairs are predicted mask confidence and IoU.

### 3.3 Mask R-CNN training

Mask R-CNN is one of the best algorithms for object instance segmentation (He et al., 2017). Intuitively, it is an extension of Faster R-CNN (Ren et al., 2015) by adding an additional branch that outputs the object mask. The results achieved by Mask R-CNN have surpassed single model results on the 2016 COCO instance segmentation competition (Lin et al., 2014). To obtain this performance, the deep neural network was trained on a huge dataset including a total of 2.5 million labeled instances in 328 thousand images. Thanks to transfer learning, rich features learned on huge datasets such as ImageNet (Deng et al., 2009) can be used in other applications by fine tuning on a new, much smaller, dataset. Using the same spirit of transfer learning, we started from the model trained on COCO by Matterport (Abdulla, 2017) and fine-tuned the network on our datasets. Particularly, we did not train all layers, and adjusted only the mask branch. We first fine-tuned the model on Dataset 1 with a learning rate of 0.005. Figure 3c shows the total loss and mask loss on the validation set during the training process using 5-fold cross-validation. The model converges around epoch 25. We term the model trained on Dataset 1, Model 1. We then tested this model on Dataset 2 which has the ground truth created by human annotators. We evaluated the model using 3 standard segmentation metrics *AP*, *AP*_75_ and *AP*_50_ (COCO detection evaluation, accessed November 17, 2021), where AP denotes average precision which is defined as the area under the precision-recall curve (PR curve). The PR curve is formed by a set of precision (the ratio of true to predicted positives) and recall (the ratio of true to actual positives) pairs. These are obtained by changing the score cutoff from 50% to 95% at a step of 5% for the intersection over union (IoU), a metric that measures the degree of overlap between a blob generated by the model and its ground truth. *AP*_75_ and *AP*_50_ are obtained when the IoU threshold is fixed at 75% and 50%.

Model 1, without training on any images from Dataset 2 i.e. without ever seeing mice in close physical proximity, achieved an *AP, AP*_75_ and *AP*_50_ of 58%, 80% and 95%, respectively as shown in Fig. 3d. Next, we investigated the network when trained on images with animals close to each other in Dataset 2. We varied the size of the training set from 20%, 40%, 60% to 80% of Dataset 2 (3 splits for all training set fractions) and fine-tuned Model 1 twelve times on them. We name the group of these twelve models, Model 1*s. The performance of the new networks gradually improves for increasing number of training samples as seen in Fig. 3d. Figure 3e shows a closer look at performance on one split for different training sizes. The set of blue lines show the kernel density estimate of *AP* achieved by Model 1* on the first splits of training sets which account for 20-80% of Dataset 2. The group of red lines show the performance of Model 1 on the same splits as above. Note that for each split, Model 1* is obtained by fine-tuning Model 1 on the other two splits. Improvements from Model 1 to Model 1* and for increasing training from Dataset 2 within Model 1* can be seen. Each row of Fig. 3f visualizes the boundary of the output blobs generated by the models (red) in comparison with human annotations (yellow). The pair of numbers for each animal are the confidence levels of the body mask that the model predicts and the IoU between the mask and its ground truth. A large improvement can be seen between the first and second images which correspond to Model 1 and Model 1* trained on 20% of Dataset 2. After that, the performance improves slowly for Model 1* trained on 40, 60 and 80% of the training set. The results show that only a small number of manually annotated frames in Dataset 2 can boost the performance significantly.

### 3.4 Tracking inference

Once masks of the animals are obtained, individual trajectories were inferred by stitching masks with the most overlapped area together. The process for this is outlined below. Here, and 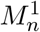 and 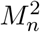 refer to the body masks returned by the algorithm and 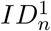 and 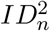 refer to animal identities that are to be assigned. IoU is the intersection over union and *n* is the frame index.

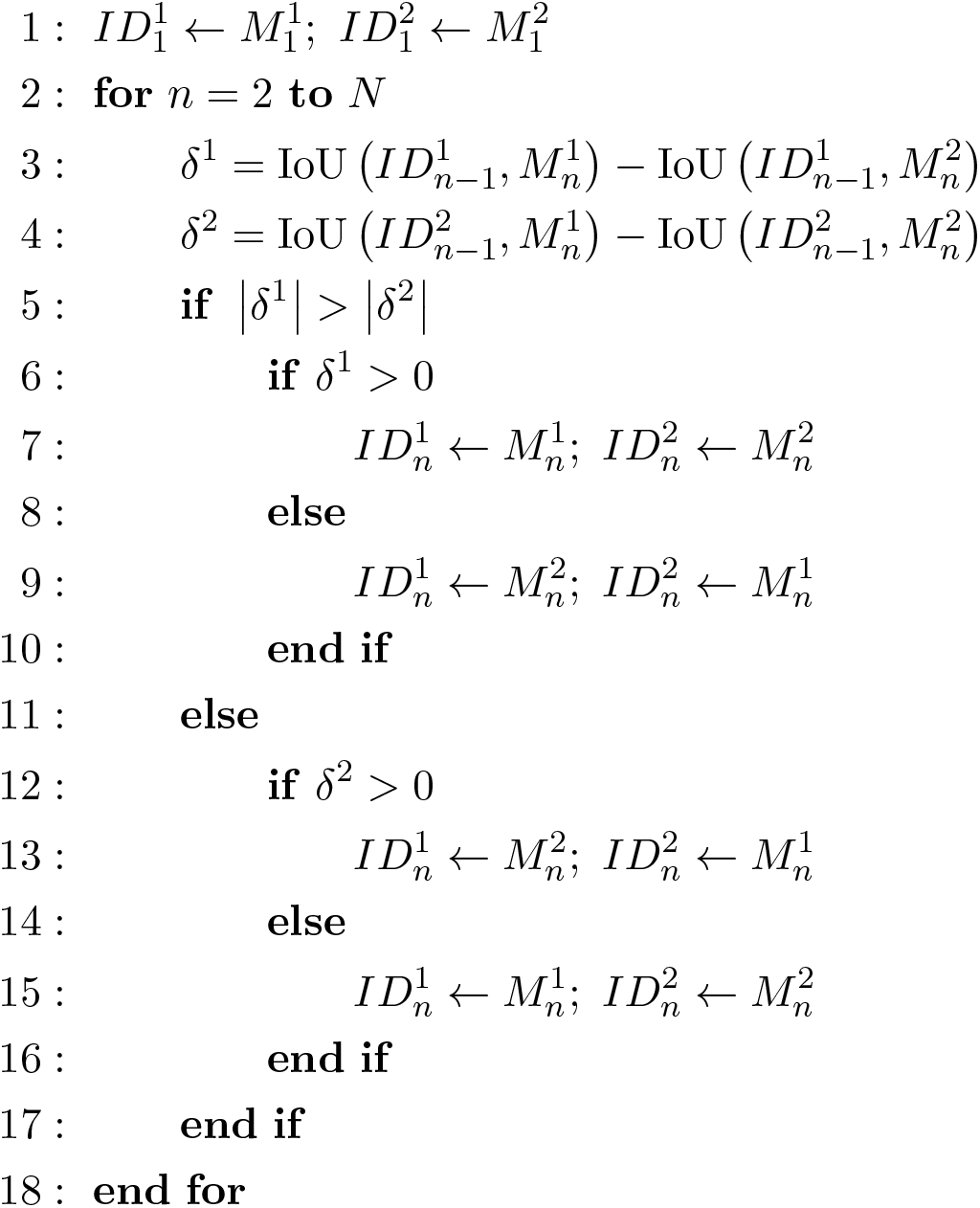

For frames that had segmentation failures, masks were replaced with the masks of the closest frame that was successfully segmented.

### 3.5 Detection of key points

We further detected key points, snout and tail-base, which are crucial for recognizing social behaviors such as ano-genital investigation and head/torso investigation. Existing approaches mainly rely on detecting users-defined key points to estimate poses before stitching them to form trajectories for each animal (Nath et al., 2019; https://sleap.ai/, accessed November 17, 2021). Such bottom-up methods work well with the visibly articulated animals such as flies and bees (https://sleap.ai/, accessed November 17, 2021). However, swapping, lost body parts and identities switches are common problems when the animals are in close contact or when body parts are occluded. This requires intensive human correction and iterative refining. To cope with these problems, we developed a hybrid method which combines deep-learningbased pose tracking and a mask-based inference. Our approach permits the incorporation of any deep learning pose estimation framework, but we chose DLC for its robust performance in comparison with others (Graving et al., 2019). For mask-based inference, we created a handcrafted detection module, termed Mask-based detection, that can assist DLC in the cases of occlusion and lost key points effectively. Also, the output of the tracking inference plays an important role in maintaining correct identities associated with detected body parts.

#### 3.5.1 Mask-based detection (MD)

This method is based on observations made in our previous work (Le et al., 2019). Specifically, the algorithm iterates over all frames and estimates the snout and tail-base location by operating on the masks to decide whether the animal’s body is parallel to the floor. This is defined by the length of the animal mask being greater than the median length over all frames and 100% overlap of the body masks with the cage floor. Otherwise, the end points are distinguished by stitching to the closest points on the previous frame.

#### 3.5.2 DeepLabCut (DLC)

A DLC model was trained only one time using the training sets described in the data preparation section above. We did not use the tracklet refining function at the inference stage. This is because we aim to develop a fully automatic tracking tool in which DLC errors would be automatically corrected by our conditional ensemble.

#### 3.5.3 Conditional ensemble

The conditional ensemble relies on the tracked body masks obtained in the tracking inference stage to output the final coordinates of snout and tail-base of each animal. Snout and tail-base coordinates generated by the deep learning framework were chosen if they were both inside an animal body mask. Otherwise, the algorithm selects the MD results.

## 4 Results

Here, were validate our approach and compare it to DLC. We also consider its generalizability to more than two mice. Finally, to demonstrate its utility and the kinds of analyses that are enabled by our approach, we quantify certain social behaviors of control or anosmic mice paired with familiar or unfamiliar mice.

### 4.1 Cross-setup validation

We evaluated the proposed algorithm on the MT dataset, specifically its performance on segmentation, identity tracking, and snout and tail-base detection.

#### 4.1.1 Segmentation

We compared the performance of the algorithm using Model 1 and the model obtained by fine tuning Model 1 on 80% of Dataset 2, which we term Model 2. The models output two body masks and confidence levels associated with each mask being a mouse. Frames where at least one confidence level was less than 0.9 or where at least one mask area was less than 20% of the largest mask area were considered as failed frames, where the two mice could not be separated. Figure 4 shows the percentage of failed frames over 12 videos in 4 categories in the MT dataset. The algorithm using Model 2 failed in less than 1% of the frames across all videos and always outperforms the algorithm using Model 1. This again confirms the previous result that training the Mask R-CNN on frames with both mice in close proximity improves the segmentation performance. Conversely, we also considered whether Dataset 1 contributed to Model 2. To answer this, we skipped training on Dataset 1, fitted a model directly on 80% of Dataset 2 and termed the resulting model Model 3. For all the categories in the MT dataset, the performance of the approach with Model 3 was not as good as the one with Model 2.

**Figure 4:**
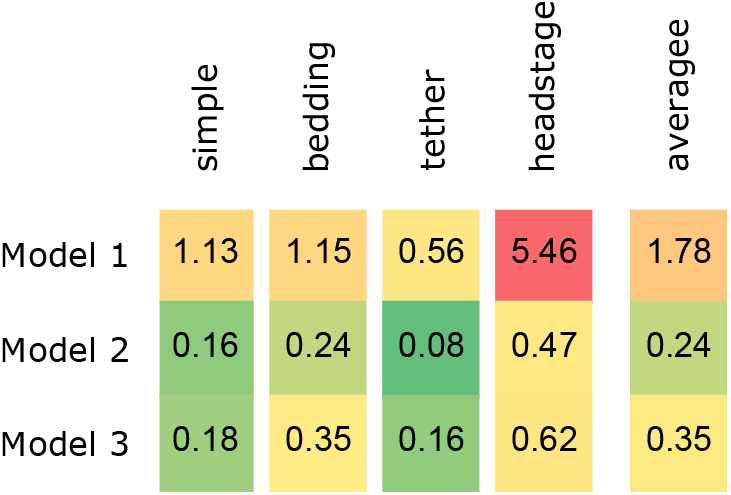
Percentage of frames with segmentation errors over 12 videos in 4 categories in the MT dataset. Model 1 is built using Dataset 1 that does not have mice in close proximity. Model 2 is a finetuned version of Model 1 incorporating manually segmented images of closely interacting mice (Dataset 2). Model 3 is trained only on Dataset 2.

#### 4.1.2 Identity tracking and key point detection

To validate the detection of key points, we manually annotated the snout and tail-base of the two mice every 10 frames in all the videos. The annotations were done by two human labelers with the split of 50:50 for each video. The coordinates of snout and tail-base obtained by MD, DLC and ensemble were compared with the human generated labels. For the most part, we followed a previously described evaluation protocol (Iqbal et al., 2017) to validate multi-animal pose tracking. To evaluate whether a body part was located correctly, we adopted the widely used PCKh (head-normalized probability of correct keypoint) metric (Zhang et al., 2019), which considers a key point to be detected correctly if it is within a certain threshold from the human annotated location. Since our videos were recorded with a fixed field of view, we apply a fixed threshold for all the animal across videos. Accordingly, we measured the length of the head as the distance from the snout to the midpoint of the shoulders in 212 randomly selected images from Dataset 2 and set the threshold to be 50% of the average head length. In addition, we evaluated center points, defined as the mid-point between the snout and tail-base, which can represent animal identity. We considered both key points and the center point as individual targets and computed the multi-object tracking accuracy (MOTA) (Bernardin and Stiefelhagen, 2008) which is widely used for evaluating multi-target tracking (Iqbal et al., 2017; Milan et al., 2016). This metric is derived from three types of error counts: missed targets (*m_n_*), the number of instances where a target is not located or is too far from the actual location; false positives (*f_pn_*), the number of instances where a target is located, but more than a threshold distance away; and mismatches (*mme_n_*), the number of instances where targets are identified but assigned incorrect identities.

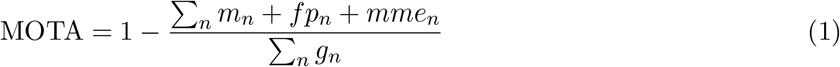

*g* is the number of targets and the subscript *n* indicates the frame number.

The upper panel of Fig. 5 shows MOTA calculated for the snout, tail-base and center using the pymotmetrics framework (https://github.com/cheind/pymotmetrics, accessed November 17, 2021). The number of identity switches that occurred in each video after analyzing with the different methods is shown in the bottom panel. On average, MOTA increases by 22% over DLC using our approach and ensembling the two has an improvement of 27%. Instances of switched identities is reduced by nearly two orders of magnitude. The worst performance, both in terms of MOTA and the number of switched identities, is seen in the headstage group of videos, particularly for the snout due to frequent occlusions caused by the headstage.

**Figure 5:**
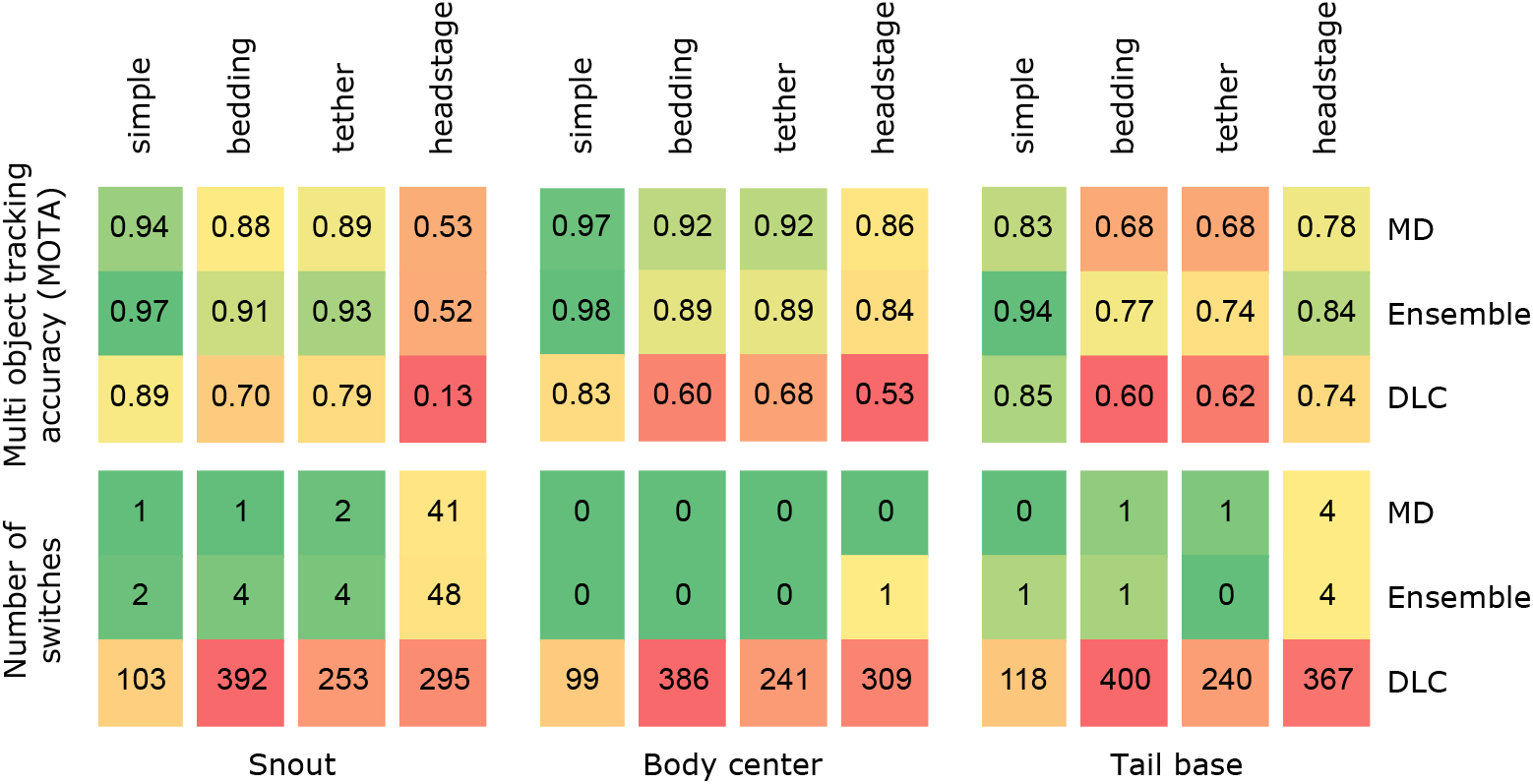
Comparison of our approaches - MD and ensemble - with DLC. The upper panel shows average Multi Object Tracking Accuracy (MOTA) and the lower panel shows total instances of switched identities across all 12 videos in the 4 categories.

Figure 6 illustrates average algorithm performance on the four groups of videos in the MT dataset, specifically regarding error distances from the ground truth. The left panel shows the fraction of frames within a certain error distance threshold. Our approach outperforms DLC in all cases. Ensembling worsens the performance somewhat in all but the headstage group, however its benefits can be seen in the right panel which shows a distribution of errors across all frames. While the median errors are a bit higher, all groups show the effectiveness of the conditional ensemble in reducing the number and size of outliers, thereby improving the overall performance. As seen earlier, the worst performance is in the headstage group especially in the detection of the snout. Results for individual videos are shown in Supplementary Figures S2-S4.

**Figure 6:**
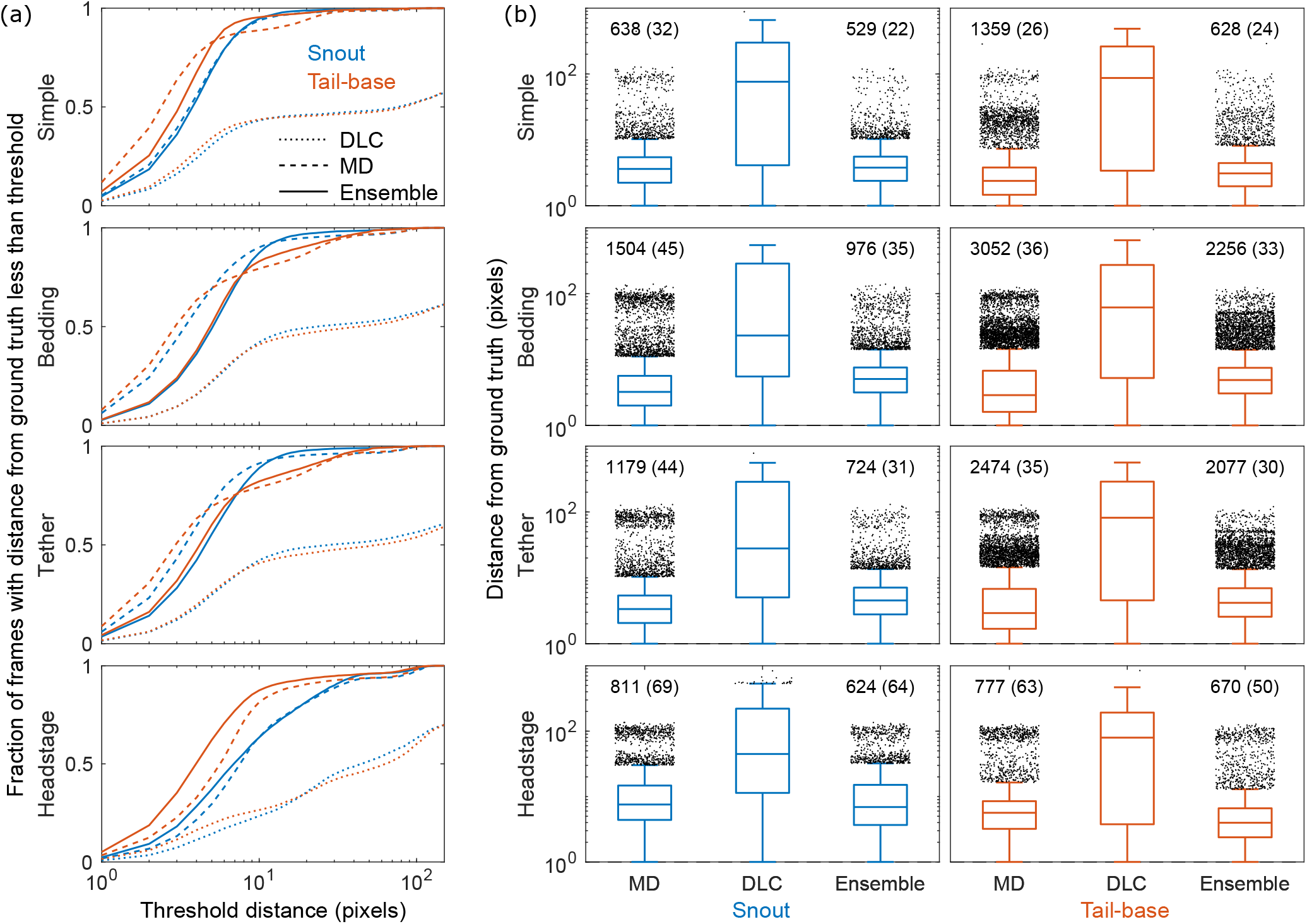
Performance across 12 videos in 4 categories. (a) Fraction of frames with the mean distance between model predictions and human annotations below a varying threshold. (b) Boxplots showing errors in MD, DLC and Ensemble models. Plots show median, 25th and 75th percentile and outliers defined as > 75th percentile + 1.5 times the inter-quartile range. Text above outliers show number of outliers and average outlier within parenthesis.

### 4.2 Generalization

We expanded the cross-setup validation to include an experiment with more than two mice. We recorded a video with 3 BTBR mice interacting for one minute (1800 frames). Fig. 7a shows some examples of good performance of the approach when coping with occlusions, while some errors can be seen in the frames shown in Fig. 7b. Figure 7c shows the trajectories of the snout and the tail-base of the mice detected by the algorithm using Model 2 trained as described above. The figure also shows the distribution of the frame-to-frame movements in x and y coordinates. The absence of large movements suggests continuity in trajectories and accurate tracking.

**Figure 7:**
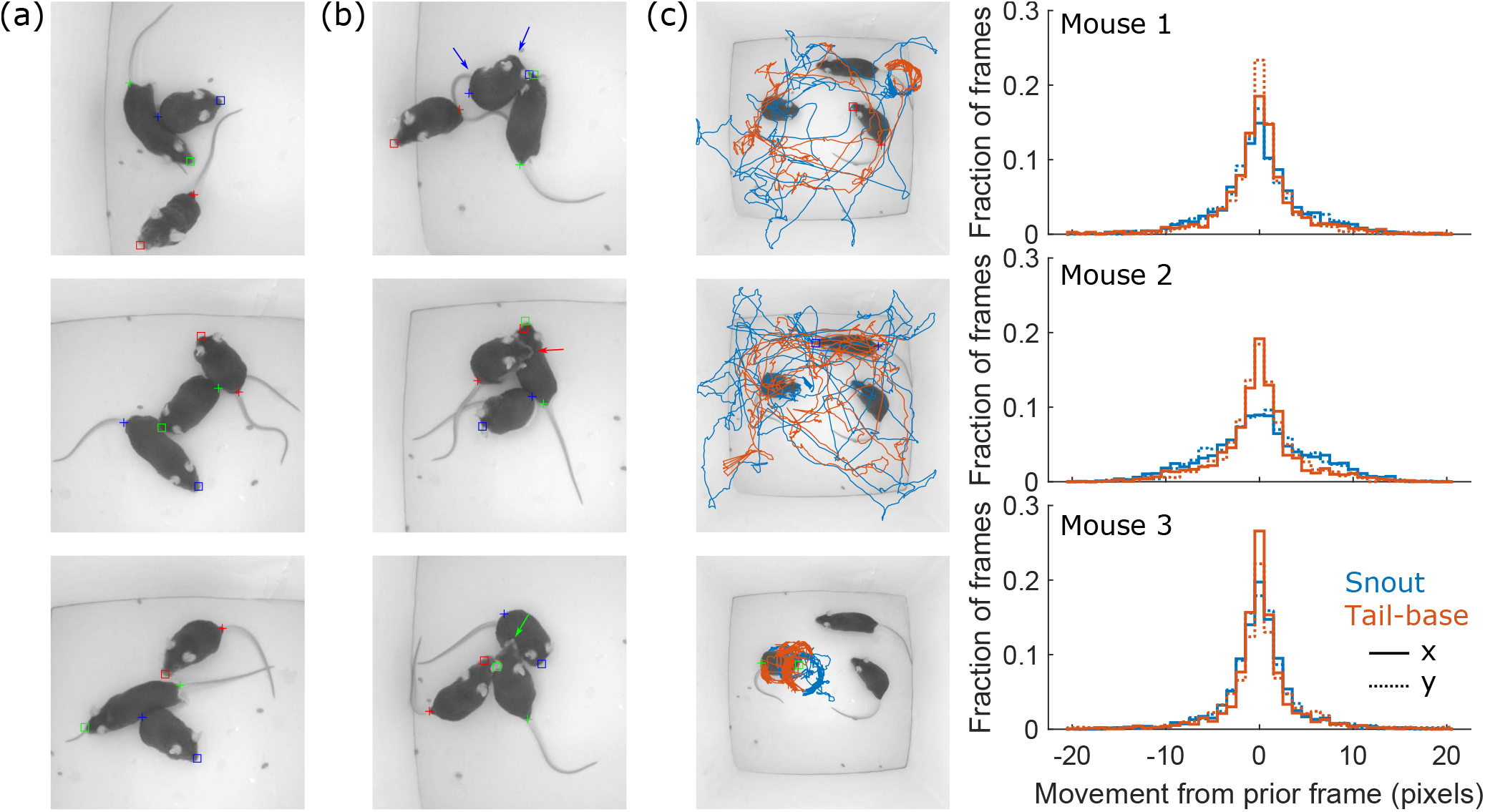
Examples of (a) good and (b) visibly inaccurate key point detection in a group of three mice. (c) Snout and tail-base trajectories for each animal along with a distribution of frame-to-frame movement in x and y coordinates.

### 4.3 Application

To test the application of our method, we used it to address a biological question: how does loss of olfaction, through ablation of the main olfactory epithelium, impact social recognition behaviors of mice. Crh-IRES-Cre mice were housed in same sex pairs. One mouse from each pair received 10 *μ*L saline (control) or 1% ZnSO_4_ (anosmic model) into each nostril to ablate the main olfactory epithelium (Ducray et al., 2002), while the partner was untreated (Fig. 8a). The next day, control or anosmic experimental mice and their partners were habituated to the tracking arena. The experimental mouse was placed into the arena first for 5 minutes before the partner was added into the arena for an additional 5 minutes. The following day (48 hrs post ZnSO_4_ or saline treatment), control or anosmic mice were returned to the tracking arena for a social recognition assay (Fig. 8b). The control or anosmic mouse acted as the “observer” (obs) and was placed into the arena first. After 5 minutes alone, the familiar partner, acting as a familiar “demonstrator” (dem) was placed in the arena. After 5 minutes, the familiar demonstrator was removed. The observer was alone for another 5 minutes before an unfamiliar demonstrator (same-sex, adult) was placed in the arena for 5 minutes. The arena was cleaned with 70% ethanol between testing observers.

**Figure 8:**
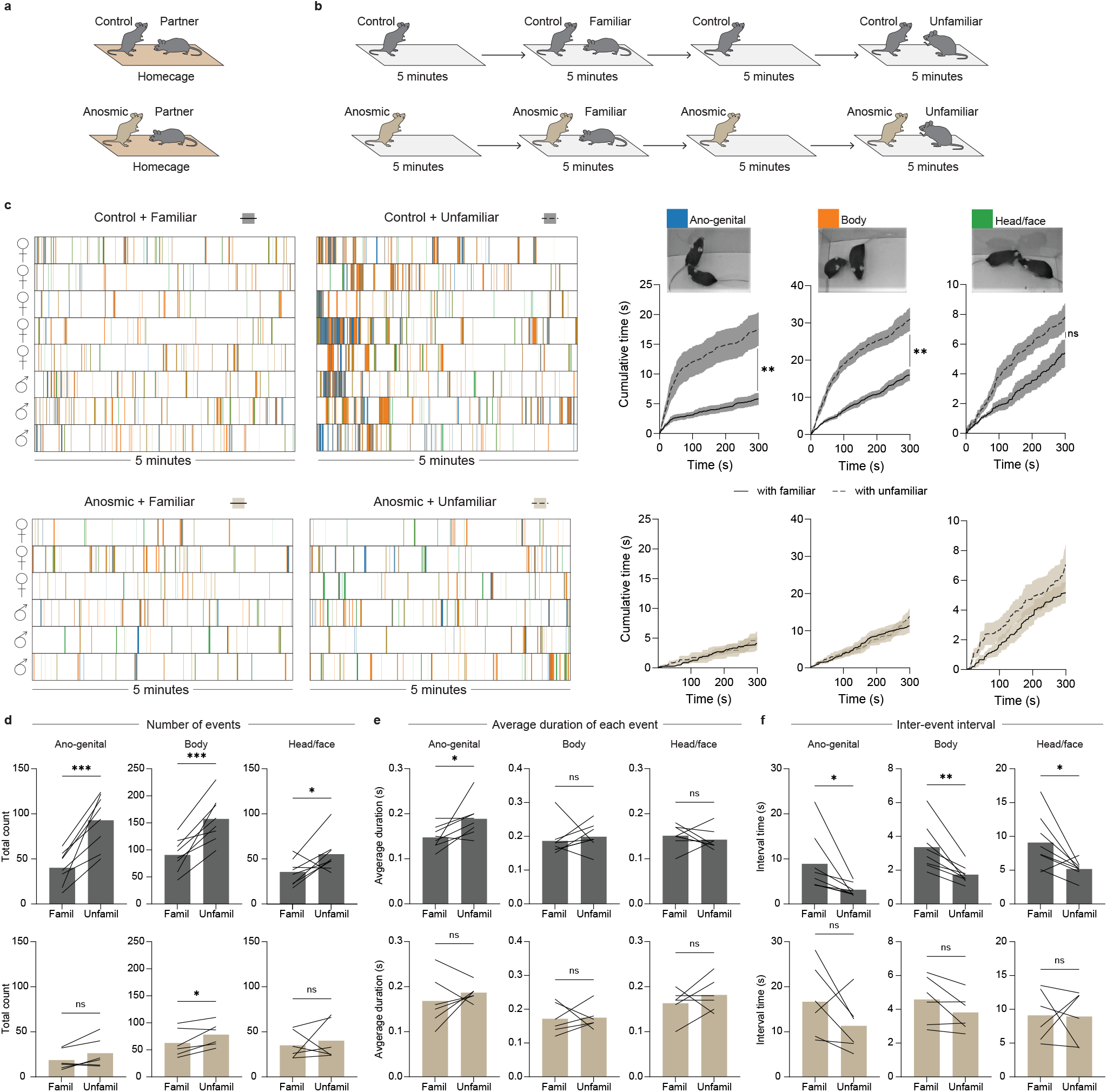
Application of the markerless mouse tracking approach showing that loss of olfaction through ablation of the main olfactory epithelium impairs social recognition behaviors in mice. (a) Mice were housed in same-sex pairs; one mouse per pair was a control (intranasal saline-treated) or anosmic (intranasal ZnSO_4_-treated). (b) Experimental paradigm for social recognition test: Anosmic/control mice acted as “observers” (Obs) and were presented with either a familiar demonstrator (Dem) or an unfamiliar demonstrator. (c) Left: Behavioral ethograms of control (top) and anosmic (bottom) observers while interacting with a familiar or unfamiliar demonstrator. Each color indicates when the snout of the observer was directed towards the ano-genital (blue), body (orange), or head/face (green) region of the demonstrator. Right: Cumulative distributions of each of the social investigative behaviors for control (top) and anosmic (bottom) observers towards familiar (solid line) or unfamiliar (dotted line) demonstrators. (d) Total number of social investigation events by control (top) and anosmic (bottom) observers directed towards the ano-genital, body, or head/face region of familiar versus unfamiliar demonstrators. (e) Average duration of each social investigation event. (f) Interval between social investigation events. *p<0.05; **p<0.01; * * *p<0.001. Paired t-test comparing social investigation towards a familiar versus unfamiliar demonstrator.

The coordinates of each mouse’s snout and tail-base, as well as the body mask, were used to determine social investigation of each mouse towards the other. Specifically, the time the snout of each mouse was directed towards the ano-genital, body, or head/face region of the other mouse was quantified (Fig. 8 & Supplementary Fig. S5). Ethograms of these 3 different types of snout-directed social behaviors reveal that that although saline treated observers investigate both familiar and unfamiliar demonstrators, there is significantly more investigation if the demonstrator is unfamiliar (Fig. 8c). In both situations, the majority of social investigation occurs during the first minute (Fig. 8c). In contrast, anosmic observers spend less time investigating their partners and do not show any preference for the unfamiliar demonstrator compared to the familiar demonstrator (Fig. 8c). Cumulative distribution graphs of time spent in each of these behaviors confirm that snout-to-ano-genital and snout-to-body investigation is increased in control observers when the demonstrator is unfamiliar (Fig. 8c). This is not true for anosmic observers (Fig. 8c). From the tracked data, we can also determine the number of social investigation events (Fig. 8d), the average duration of each event (Fig. 8e), and the average time between events (Fig. 8f). In control observers, the number of social investigation events increased (Fig. 8d) and the inter-event interval decreased (Fig. 8f), when the demonstrator was unfamiliar, for all types of investigation (anogenital, body, head/face). The average duration of only ano-genital investigation was increased when the demonstrator was unfamiliar (Fig. 8e). For anosmic observers, the number of body-directed investigation events increased when the demonstrator was unfamiliar (Fig. 8d), while no other parameters were significantly changed. These findings suggest that intact olfaction is required for many social behaviors that demonstrate social discrimination of a familiar versus unfamiliar conspecific.

Since both animals are tracked, we can analyze demonstrator mouse behavior as well. Demonstrator mice show very little snout-directed social investigation of observer mice. Time spent engaged in these different types of investigation are also not altered if the demonstrator is familiar or unfamiliar to the observer (Supplementary Fig. S5). Thus, social investigation behaviors appear to be uni-directional in this case, exhibited predominantly by the mouse that was in the arena first, i.e. the observer.

Since our approach generates body masks in addition to the snout and tail-base coordinates, we could also determine how much time observers and demonstrators spent in contact with one another by detecting at what times the masks overlapped. By removing the snout-directed social investigation times from this data, we could determine the amount of time each pair spent touching that was not because of observer-to-demonstrator or demonstrator-to-observer snout-directed investigation. Behavioral ethograms indicate that pairs of mice in which one is anosmic spend more time in contact compared to control pairs (Fig. 9a).

**Figure 9:**
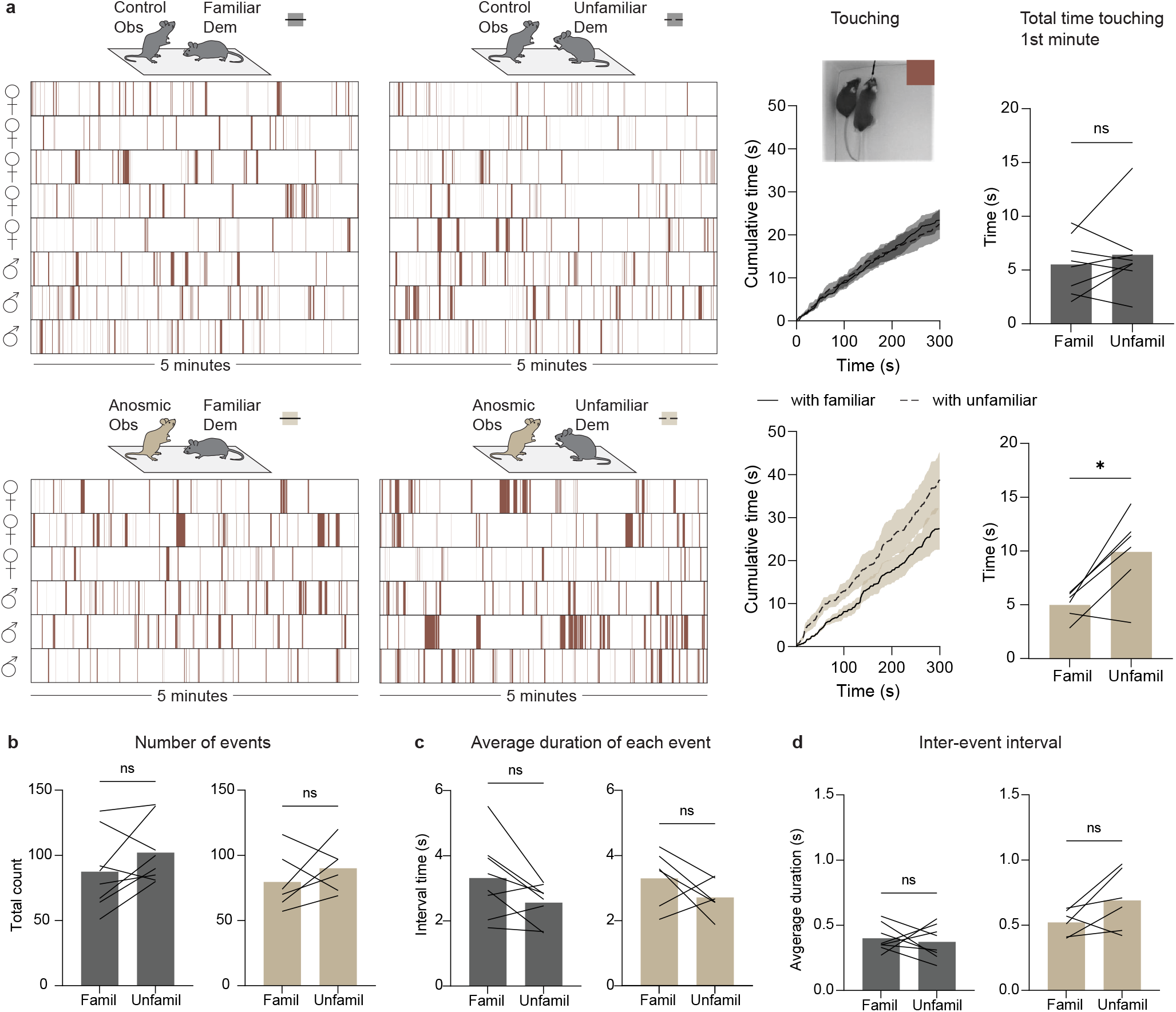
Application of the markerless mouse tracking approach showing that anosmic mice spend more time in contact (non-snout-directed contact) with unfamiliar versus familiar demonstrators. (a) Left: Behavioral ethograms of control (top) and anosmic (bottom) observers while interacting with a familiar or unfamiliar demonstrator. Each brown bar indicates when the mice were in contact (touching) that was not as a result of snout-directed investigation by the observer or demonstrator. Right: Cumulative distributions of touching in pairs of control (top) or anosmic (bottom) observers with familiar (solid line) or unfamiliar (dotted line) demonstrators. Bar graph shows total time spent touching during the 1st minute of the 5-minute period. (b) Total number of touching events between control (left) or anosmic (right) observers, with familiar or unfamiliar demonstrators. (c) The average duration of each touching event. (d) The inter-event interval between touching events. *p<0.05. Paired t-test comparing touching between observers and familiar versus unfamiliar demonstrators.

Plotting these data as cumulative distributions shows that anosmic observers spend more time in contact with unfamiliar demonstrators than familiar demonstrators, but only near the beginning of the 5-minute period (Fig. 9a). Taking the average time spent touching during the first minute shows that anosmic observers do spend more time in contact with unfamiliar demonstrators than familiar demonstrators (Fig. 9a). The number of contact events (Fig. 9b), average duration of events (Fig. 9c), and inter-event intervals (Fig. 9d) for the total 5-minute period were not altered by the familiarity of the demonstrator. These findings show that anosmic mice might behaviorally discriminate between unfamiliar and familiar conspecifics through changes in bodily contact rather than snout-directed investigation.

Tracking the centre point of observers and demonstrators shows that when the observer is anosmic, the pair travel a shorter distance than control pairs (Fig. 10a). Quantifying the distance covered shows that anosmic observers move less than their non-anosmic demonstrators (Fig. 10b), suggesting that loss of olfaction impacts exploratory behaviors in general. Anosmic observers also spend more time stationary when they are in the presence of an unfamiliar demonstrator versus a familiar demonstrator (Fig. 10c). This again shows behavioral discrimination of an unfamiliar versus familiar conspecific by anosmic mice that is independent of snout-directed behaviors. Looking at velocity of observer and demonstrator mice in 1s bins over the 5-minute period shows that the velocity of control observers and their demonstrators tend to be correlated (75% of pairs correlated, Supplementary Table S2; example shown in Fig. 10d). Velocity of pairs where the observer is anosmic tend to not be correlated (75% of pairs not correlated, Supplementary Table S2; example shown in Fig. 10d). This suggests that intact olfaction is required for coordinated behaviors in a pair of mice.

**Figure 10:**
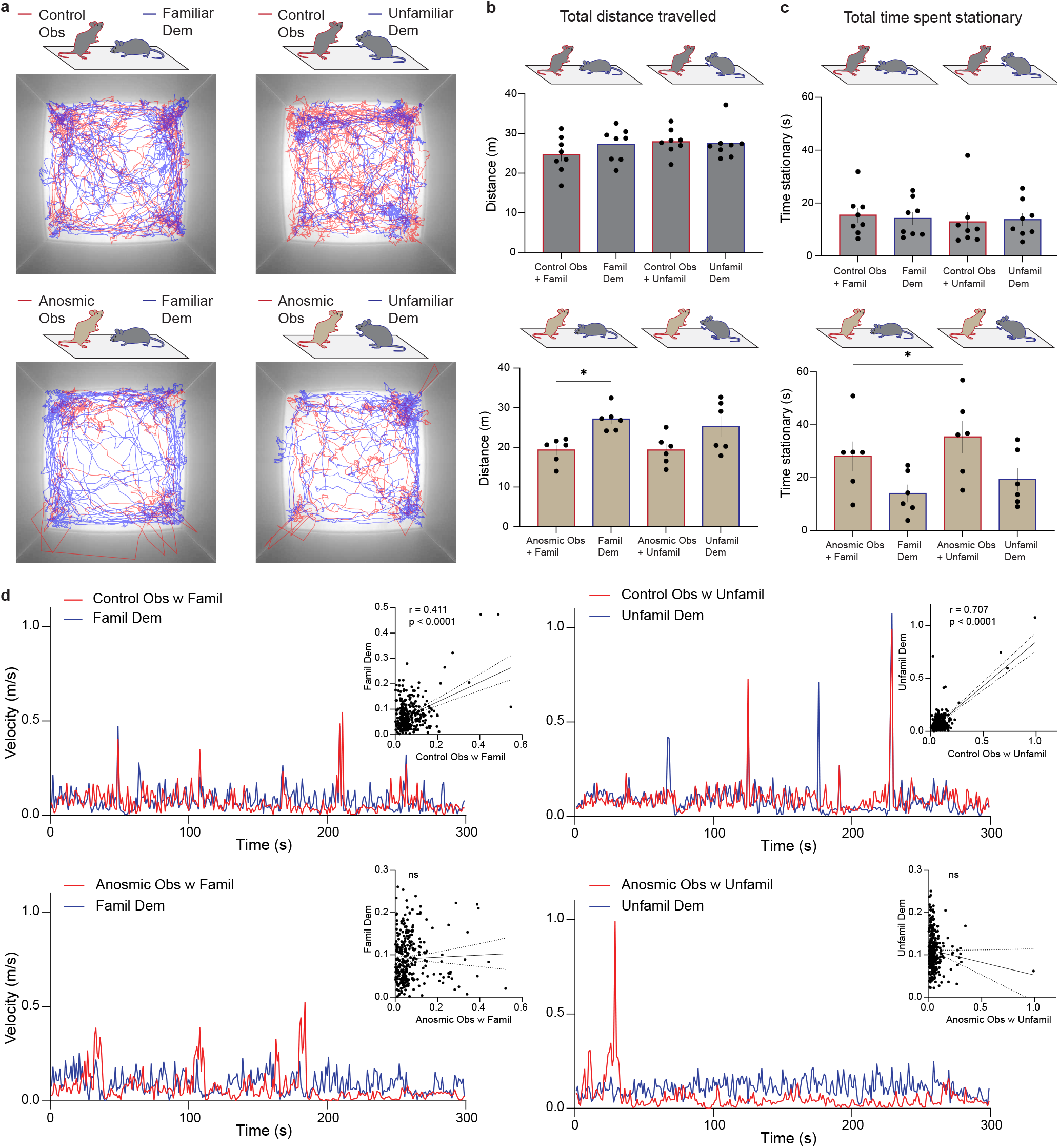
Application of the markerless mouse tracking approach showing distance travelled and velocities of pairs of mice (observer + demonstrator). (a) An example of raw tracking data of observers (red) and demonstrators (blue), where observers were control (top) or anosmic (bottom) with familiar (left) or unfamiliar demonstrators (right). (b) Total distance travelled by observers and their familiar or unfamiliar demonstrators where the observer was control (top) or anosmic (bottom). (c) Total time spent stationary by observers and their familiar or unfamiliar demonstrators where the observer was control (top) or anosmic (bottom). (d) Examples of velocities (1 s bins) of observers (red) and demonstrators (blue) with familiar or unfamiliar demonstrators where the observer was control (top) or anosmic (bottom). Inset graphs show Pearson correlation analyses of observer velocities versus demonstrator velocities. (b) and (c) *p<0.05, One-way ANOVA with Sidak’s multiple comparisons test.

## 5 Discussion

Reliably tracking multiple animals is critical for automated behavioral assays evaluating social interactions. However, it is extremely time-consuming to evaluate the quality of animal tracking systematically, because the ground truth needs to be generated manually. As of now, there are no publicly available benchmark datasets, and only a few attempts have been made to evaluate tracking methods (Lorbach et al., 2015; Salem et al., 2015). Some reports have indicated that behavioral assays in close-to-natural arenas are more ethologically relevant than those in bare cages (Castelhano-Carlos et al., 2014; Puścian et al., 2016). Additionally, advances in optogenetics, fiber photometry and tethered or wireless electrophysiology permit simultaneous physiological manipulation and recording along with behavioral tests. These require animal-mounted headstages and fiberoptic or electrical cables. Thus, tracking algorithms that are robust to the presence of bedding and other enrichments as well as various implants would be particularly useful.

Here we developed an approach that combines conventional tracking and a deep learning-based instance segmentation method that advances tracking of social behaviors in two important ways: First, we decreased the computational cost of deep neural networks that are sufficiently powerful to cope with challenges like social contact between animals to extract animals’ masks but must operate on a frame by frame basis. We took advantage of good contrast and used conventional methods to obtain blobs when the animals were apart and only applied the Mask R-CNN in cases when the animals were interacting. This also reduced the required training set size for the Mask R-CNN since the mouse body is highly deformable and can result in a great number of postures. Second, the conventional method can be implemented in parallel for a batch of frames using multiple CPU cores. Consequently the hybrid approach is much more time efficient compared to the approach using only Mask R-CNN. In addition, the ensemble results of key point detection have shown that combining handcrafted inference with the deep neural network can improve the performance of the system considerably, and that the method remains robust with different settings and implants. This suggests that injecting human knowledge can be very helpful to mitigate the limitations of end-to-end approaches. The videos in the mouse tracking database and an implementation of our method are available at https://github.com/MaSoMoTr/MaSoMoTr for the research community to further develop tracking and key point detection for automated analysis of social behaviors.

Current approaches directed to quantifying social behavior have unique advantages and limitations. Researchers need to carefully consider these factors to determine the suitable methods for the unique nature of any experiment. If a reasonable contrast between the background and the animals is maintained, the background appearance can be modeled at every pixel location, and then the foreground can be segmented (Noldus et al., 2002; Branson et al., 2009; Swierczek et al., 2011). This approach, however, is largely limited to individual animals since tracking multiple animals in the same arena is prone to occlusion or “identity switching” because of frequent social interactions at close range.

As mentioned earlier, approaches that use RFID can provide accurate location tracking of multiple animals but do not provide information on the positions of important body regions. RFID methods, requiring surgically implanted RFID tags in the animals, have been used to track individuals in group cages (de Chaumont et al., 2019; Weissbrod et al., 2013; Schaefer and Claridge-Chang, 2012; Galsworthy et al., 2005). This allows for experiments in semi-natural and less controlled environments (Weissbrod et al., 2013) with accessories such as feeders, bridges, and shelters. Besides requiring surgery, another limitation is that integration of detection coils might complicate the testing arena (Lewejohann et al., 2009). Animals also need to recover for a few days post-surgery, which imposes constraints on experimental designs and timelines.

Other approaches have used physical marking of individual animals to assist automatic video-based tracking systems. Human hair bleach has been used to create unique patterns on mouse fur, enabling multi-day tracking (Ohayon et al., 2013). Noldus EthoVision XT 12 (Lorbach et al., 2017), a commercial system, tracks white rats and their body parts after marking animals with a black marker. Although such marking approaches can make tracking algorithms more robust and avoid switching identities, they have drawbacks (Walker et al., 2013). Markings may fade over time requiring reapplication under restraint and handling by humans which is time-consuming and stressful for the animals (Burn et al., 2008; Hurst and West, 2010). Certain compounds in marking agents may also alter behavior in some assays (Burn et al., 2008; Hershey et al., 2018). Furthermore, marking procedures must be determined in advance, and implemented carefully and consistently.

Several markerless tracking methods have been proposed to track animals. One example is carrying out the experiments with a pair of back and white mice (Hong et al., 2015; Burgos-Artizzu et al., 2012). However, this approach is not very broadly applicable since commonly used genetically modified animals are often dark colored (The Jackson Laboratory, accessed November 17, 2021), and littermate controls are often of the same fur color as the experimental group.

When multiple animals are involved, social contacts routinely lead to occlusions, causing identity errors. To cope with occlusion, Matsumoto proposed utilizing multiple 3D depth cameras with different viewpoints to reconstruct 3D positions of rats’ body parts in order to analyze social and sexual interactions (Matsumoto et al., 2013). While a depth camera provides additional information about posture and body shape compared to a standard camera, the technique requires a relatively complex setup and the process is computationally intensive. Unger et al. (Unger et al., 2017) introduced an unsupervised-learning method to track the animals in an uncluttered setting by learning the shape information of each animal, and then segmenting the animal by matching the current shape with corresponding catalog. The method needs considerable preprocessing including human annotation as the shape catalog has to be built for each animal used in an experiment, and can not be reused for any other animal.

Convolutional Neural Network based trackers have achieve state-of-the-art performance on benchmark datasets (Smeulders et al., 2013; Wu et al., 2015) and have been applied to pose estimation of mice housed in laboratory cages (Mathis et al., 2018; Pereira et al., 2019; Graving et al., 2019). DeepLabCut version 2.2 (Nath et al., 2019) and SLEAP (Pereira et al., 2019; https://sleap.ai/, accessed November 17, 2021) enable tracking unmarked animals. An important feature of such end-to-end deep learning models is their flexibility: with enough carefully prepared training data, virtually any experimental setting can be handled with asymptotically improving performance. For example, they can be trained to track several species and applied to track user-defined points like whiskers and paws. A major limitation of these models is that they require significant human intervention to correct mistakes such as swapped features when tracking multiple animals with limited training data, as indicated by our test on tracking two interacting mice. Additionally, performance was further reduced on videos in enriched arenas or when animals had implants. With comprehensive training data collection and curation, performance can be improved, but at the cost of significant human effort.

Our algorithm was trained using a single video in a simple, uncluttered setting. We used 1060 non-curated images with animals apart and 212 manually segmented images when the animals were closely interacting. Our results, obtained without any human intervention in videos that included bedding, and animals with fiberoptic or headstage implants, demonstrated a near elimination of switched identities and a 10% improvement in tracking accuracy over DLC. Additionally, the results for three mice show the potential to apply our approach to experiments involving a group of mice. However, our method is not as flexible as end-to-end deep learning frameworks which, for example, can be applied to other species or tracking arbitrary user-specified body parts. Instead, it was designed specifically for accurately tracking socially interacting mice, and locating their snouts and tail-bases with minimal training and human intervention. A more specific limitation of our method is that extensive occlusion caused by big implants such as EEG headstages is still a challenge. Thus, manual validation for identity assignments in tracking results is recommended to correct swapping identities in challenging situations.

By applying our markerless mouse tracking tool to address a biological question, we made several discoveries. We specifically tracked the coordinates of two mice, where one mouse was either a control or had been made anosmic. Our results indicated that while control mice show an increase in snout-directed social investigation of a conspecific when the conspecific is unfamiliar versus familiar, this is not true for anosmic mice. Given that rodents rely heavily on olfactory cues in social recognition (Sanchez-Andrade and Kendrick, 2009; Oettl et al., 2016), this result was expected. We found evidence, however, of behavioral discrimination of familiar versus unfamiliar conspecifics in pairs of mice where one was anosmic. Specifically, touching/contact that was not snout-directed investigation from either mouse was increased when the conspecific was unfamiliar versus familiar. We also showed that anosmic mice spend more time stationary when in the presence of an unfamiliar versus a familiar mouse. These subtle differences suggest that anosmic mice use non-olfactory cues to discriminate familiar and unfamiliar conspecifics. Finally, we demonstrate additional value of this tool in tracking velocities of interacting animals. We show that majority of pairs of mice display correlated velocities, indicating coordinated movements; when one mouse is anosmic, however, the pair are less likely to have correlated velocities. Such application of this tool opens doors to many new possibilities of analyses that may hold specific value depending on the research question. Acquiring data on social contact (snout-directed or other), geographic location, and distance travelled, all with temporal resolution, allows for combining of data sets to address questions like: at what velocity does snout-directed social investigation occur? Where in the arena do mice engage in social contact? Do snout-directed social investigations follow a specific pattern? Do mice show reciprocal investigation patterns after a certain time? And how do these all change in mice with a specific genetic mutation, or after an environmental perturbation? While these questions are beyond the scope of the current study, they can now readily be addressed using this tool.

## 6 Conclusion

We proposed a hybrid approach for tracking mice and their features in experiments involving two mice of the same color. We evaluated the method using cross-setup videos in two aspects: tracking identities and key point detection. The results obtained with a small number of training images and without any human correction have shown the utility of the proposed method.

## 7 Supplementary Material

**Figure S1:**
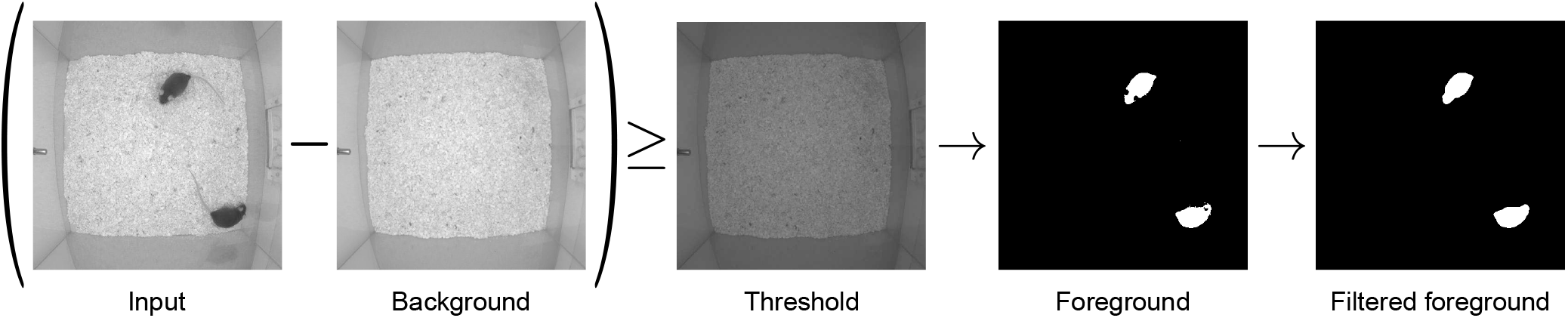
Foreground detection includes background subtraction, thresholding and a series of morphological operations: filtering, closing and opening.

**Figure S2:**
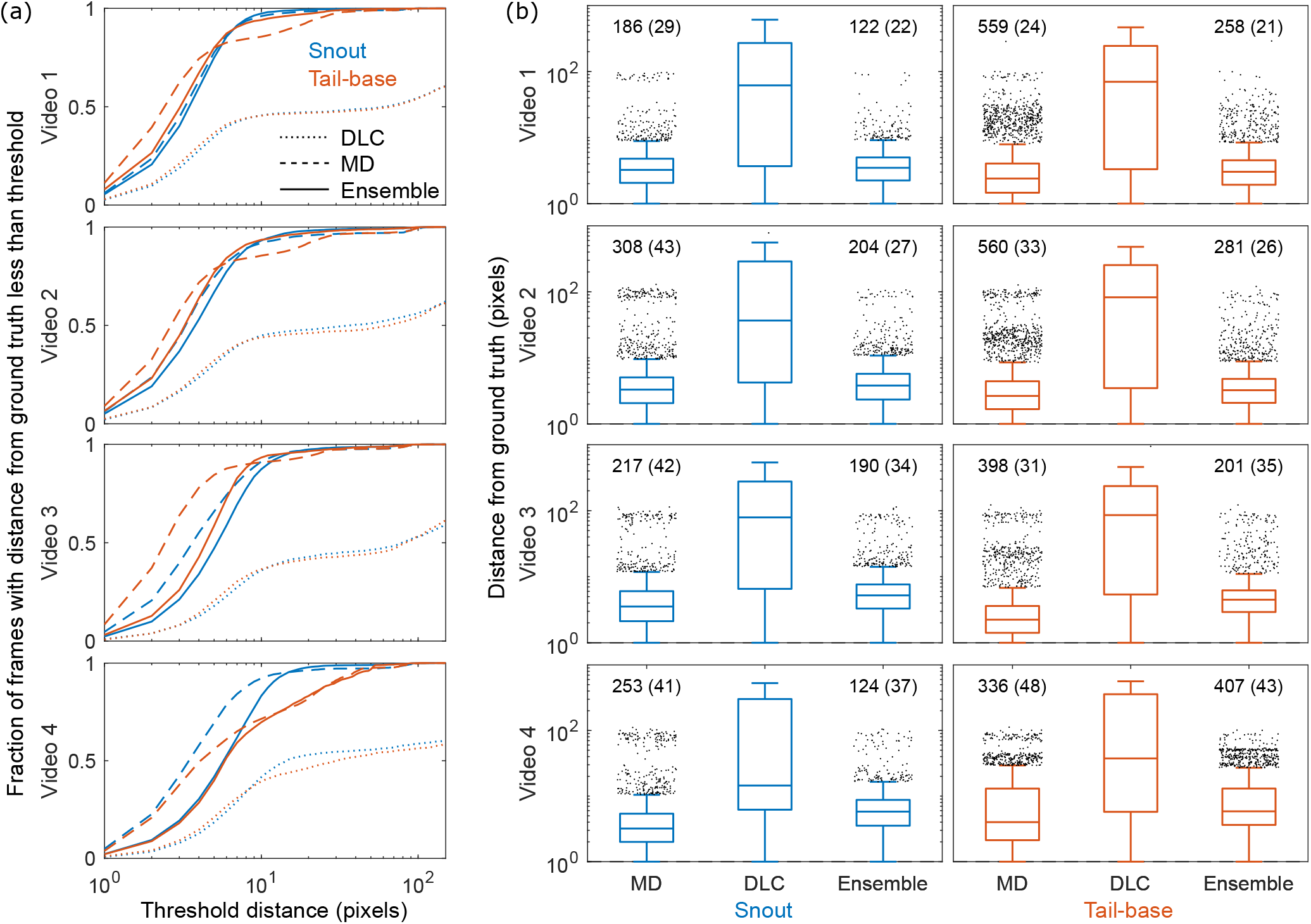
Performance across videos 1-4. (a) Fraction of frames with the mean distance between model predictions and human annotations below a varying threshold. (b) Boxplots showing errors in MD, DLC and Ensemble models. Plots show median, 25th and 75th percentile and outliers defined as > 75th percentile + 1.5 times the inter-quartile range. Text above outliers show number of outliers and average outlier within parenthesis.

**Figure S3:**
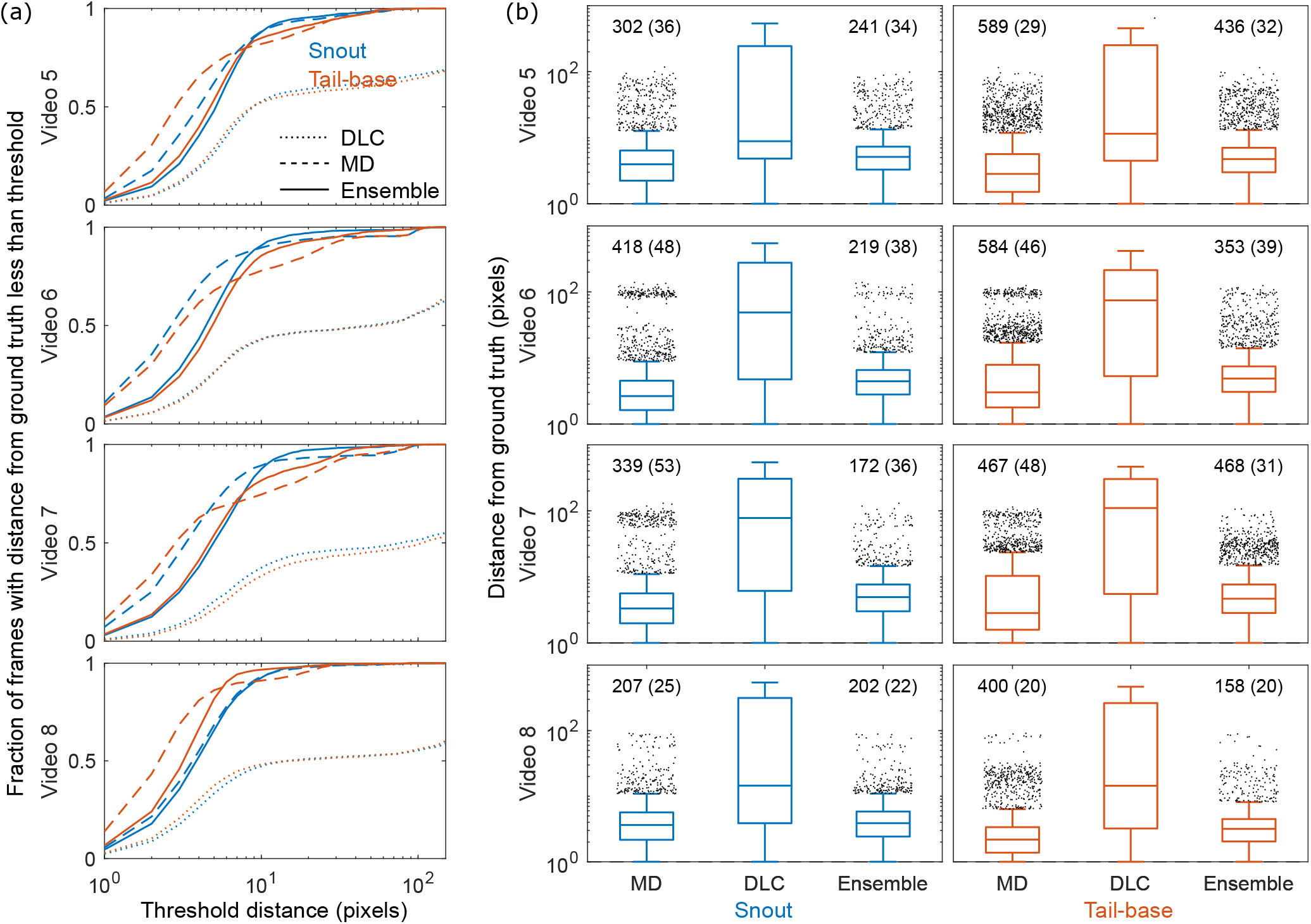
Performance across videos 5-8. (a) Fraction of frames with the mean distance between model predictions and human annotations below a varying threshold. (b) Boxplots showing errors in MD, DLC and Ensemble models. Plots show median, 25th and 75th percentile and outliers defined as > 75th percentile + 1.5 times the inter-quartile range. Text above outliers show number of outliers and average outlier within parenthesis.

**Figure S4:**
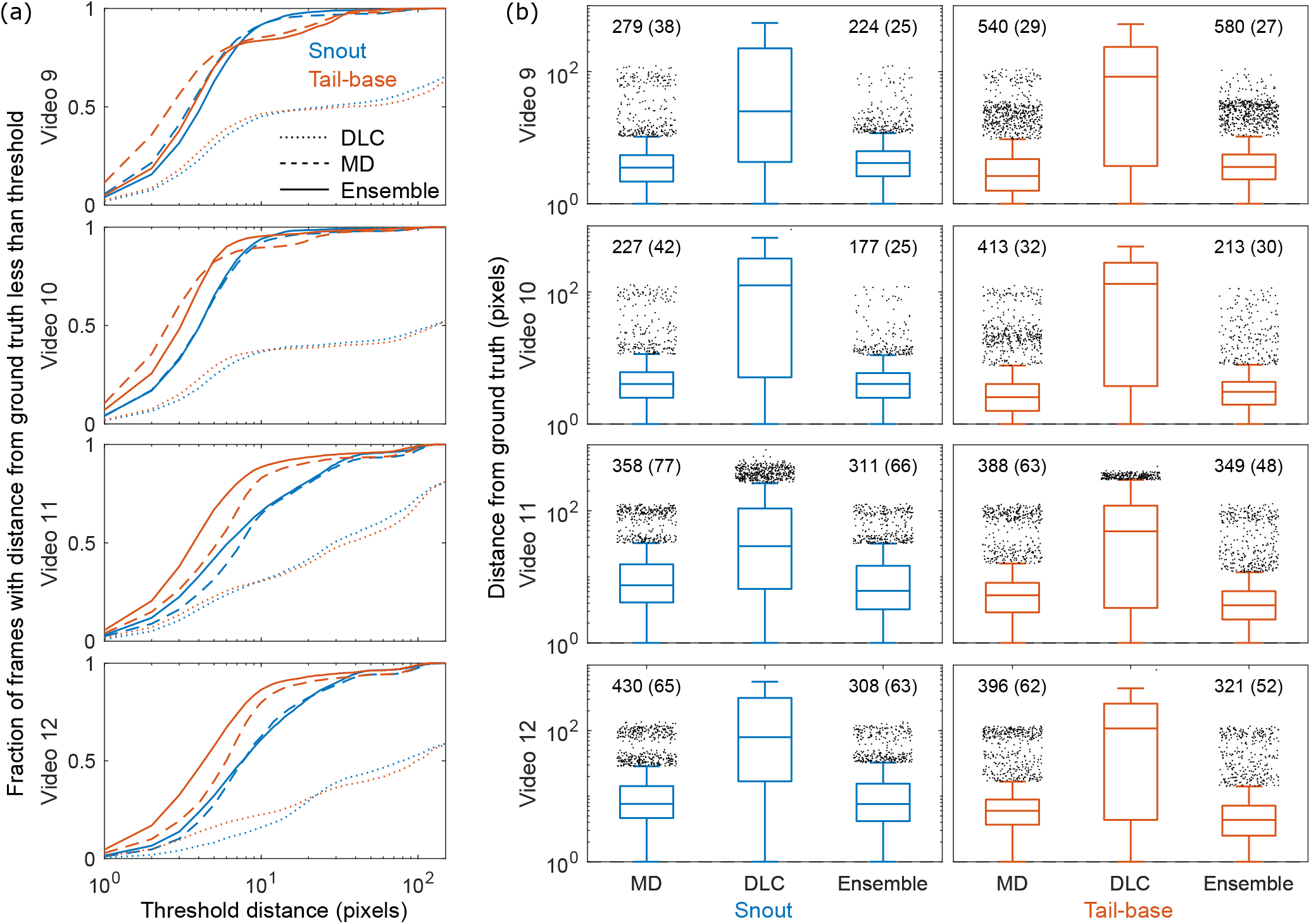
Performance across videos 9-12. (a) Fraction of frames with the mean distance between model predictions and human annotations below a varying threshold. (b) Boxplots showing errors in MD, DLC and Ensemble models. Plots show median, 25th and 75th percentile and outliers defined as > 75th percentile + 1.5 times the inter-quartile range. Text above outliers show number of outliers and average outlier within parenthesis.

**Figure S5:**
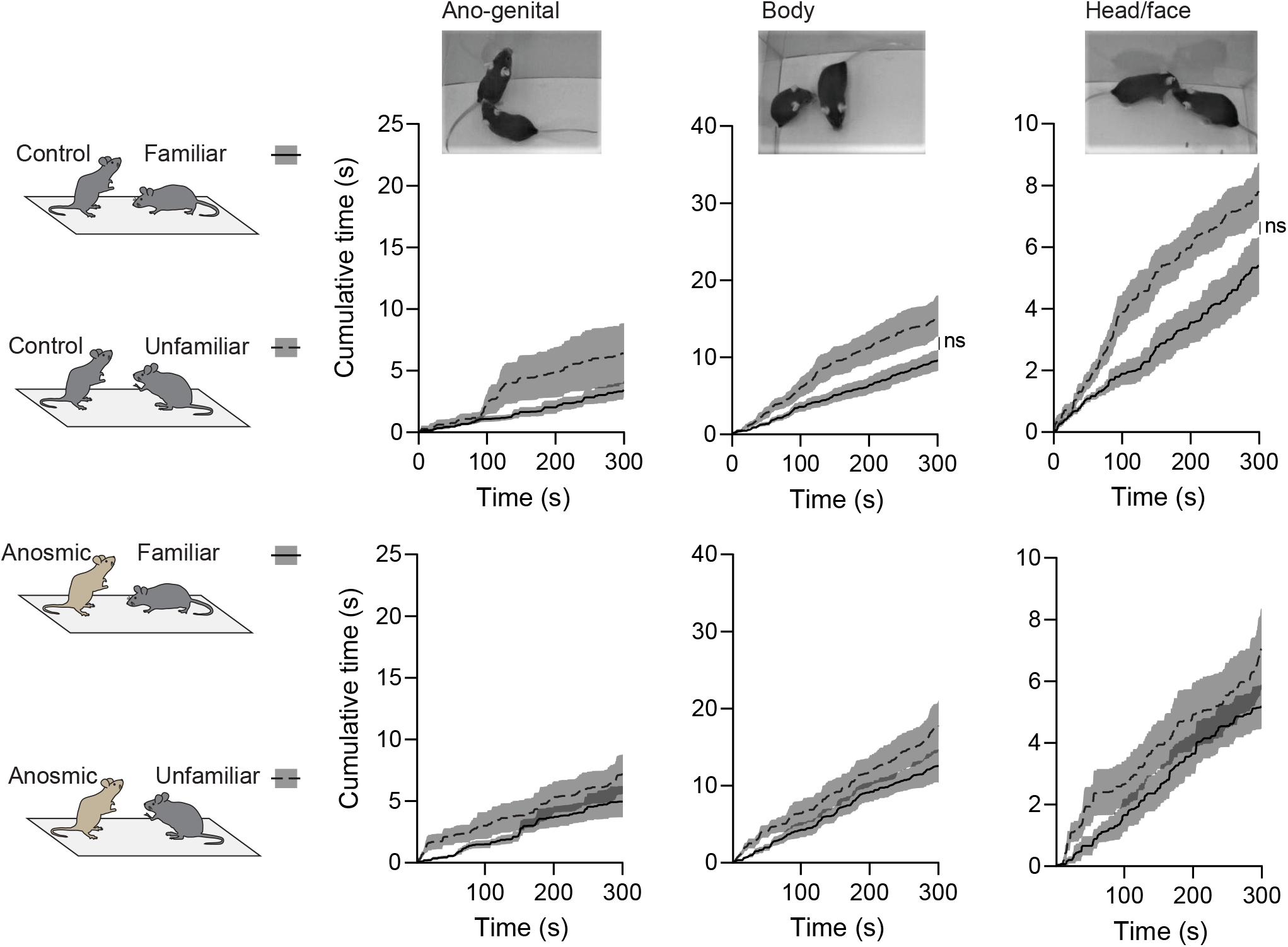
Social investigation behaviors of familiar or unfamiliar demonstrators towards control or anosmic observers. Social investigation behaviors include when the snout of the demonstrator was directed towards the ano-genital, body, or head/face region of the observer. Cumulative distributions show that familiar (solid line) and unfamiliar (dotted line) demonstrators spend similar amount of time engaged in each social investigation behaviour when with a control (top) or anosmic (bottom) observer.

**Table S2:** Velocities (calculated in 1 second bins as metres per second) correlated more frequently in control observer + demonstrator pairs (75%) than in anosmic observer + demonstrator pairs (25%)

| Treatment | Sex | Famil or Unfamil pair | Velocities correlate | r value | p value |
| --- | --- | --- | --- | --- | --- |
| Control | F | Famil | no | - | - |
| Control | F | Unfamil | yes | 0.1916 | 0.0009 |
| Control | F | Famil | no | - | - |
| Control | F | Unfamil | no | - | - |
| Control | F | Famil | yes | 0.2158 | 0.0002 |
| Control | F | Unfamil | yes | 0.1293 | 0.0254 |
| Control | F | Famil | yes | 0.411 | <0.0001 |
| Control | F | Unfamil | yes | 0.7067 | <0.0001 |
| Control | F | Famil | yes | 0.4741 | <0.0001 |
| Control | F | Unfamil | yes | 0.2865 | <0.0001 |
| Control | M | Famil | yes | 0.1969 | 0.0006 |
| Control | M | Unfamil | yes | 0.1492 | 0.0098 |
| Control | M | Famil | yes | 0.1699 | 0.0032 |
| Control | M | Unfamil | yes | 0.1848 | 0.0013 |
| Control | M | Famil | no | - | - |
| Control | M | Unfamil | yes | 0.3839 | <0.0001 |
| Anosmic | F | Famil | no | - | - |
| Anosmic | F | Unfamil | yes | 0.1207 | 0.0371 |
| Anosmic | F | Famil | no | - | - |
| Anosmic | F | Unfamil | no | - | - |
| Anosmic | F | Famil | yes | 0.4585 | <0.0001 |
| Anosmic | F | Unfamil | yes | 0.2173 | 0.0002 |
| Anosmic | M | Famil | no | - | - |
| Anosmic | M | Unfamil | no | - | - |
| Anosmic | M | Famil | no | - | - |
| Anosmic | M | Unfamil | no | - | - |
| Anosmic | M | Famil | no | - | - |
| Anosmic | M | Unfamil | no | - | - |

